# Inflammation is the Driver of Butyrate-Producing Bacteria Change in Interleukin10 Knockout Mice

**DOI:** 10.64898/2026.01.15.699743

**Authors:** Ji Yeon Kim, Bohye Park, Olivia F. Riffey, Madison L. Bunch, Jeremiah G. Johnson, Zachary M. Burcham, Dallas R. Donohoe

## Abstract

**Background:** Alterations of gut microbiota have been implicated in the development of inflammatory bowel disease. Specifically, patients with IBD show the reduced levels of gut bacteria to produce butyrate, a crucial metabolite for maintaining gut homeostasis, along with decreased levels of fecal butyrate. However, there is limited research on changes in butyrate-producing bacteria at various taxonomic levels during the development of inflammatory bowel disease.

**Results:** We investigated the changes of butyrate-producing bacteria in interleukin10 knockout mice, a suitable IBD model, as these mice require gut microbiota to develop spontaneous chronic colitis. Our findings indicate increased inflammation and a metabolic shift from butyrate oxidation toward glycolysis in 9-week-old interleukin10 knockout mice. Furthermore, we observed significant changes in two terminal enzymes involved in butyrate production: a significant increase of butyrate kinase and a significant decrease of butyryl-CoA:acetate-CoA-transferase. These observations align with an increased abundance of *Coprococcus comes*, which utilizes butyrate kinase, and a decreased abundance of *Faecalibacterium prausnitzii* that utilizes butyryl-CoA:acetate-CoA-transferase. Moreover, reduced levels of acetate, a necessary co-substrate for butyryl-CoA:acetate-CoA-transferase activity, were observed in interleukin10 knockout mice.

**Conclusions:** These findings enhance our understanding of changes in butyrate-producing bacteria populations at various taxonomic levels, ranging from phylum to gene level in 9-week-old interleukin10 knockout mice. Furthermore, these data suggest a potential for diagnosing IBD at an early stage by analyzing the composition of butyrate-producing bacteria.

## Background

Inflammatory bowel disease (IBD) including Crohn’s disease (CD) and ulcerative colitis (UC) is a gastrointestinal disease characterized with a chronic and recurring inflammation. Although IBD’s precise pathogenesis remains unclear, extensive evidence supports the pivotal role of gut microbiota in promoting IBD pathogenesis [1, 2]. This is evidenced by studies on antibiotic treatment of IBD and experimental colitis models [3–5]. As a result, dysbiosis, an imbalance or disruption of the intestinal microbiota, emerges as a common feature of IBD. Several studies have reported the changes in the composition of gut microbiota in IBD patients, characterized by an increased abundance of Proteobacteria and Bacteroidetes and a decrease in Firmicutes [6–10].

Butyrate, a short-chain fatty acid (SCFA), is produced through anaerobic microbial metabolism in the gastrointestinal tract. It is the primary energy source for the colonic epithelial cells and contributes to the maintenance of intestinal homeostasis, including modulating immune response to protect the intestinal barrier function [11–13]. Butyrate-producing bacteria synthesize butyrate from butyryl CoA using two different enzymes, butyrate kinase and butyryl-CoA:acetate-CoA-transferase (butyrate transferase) [14, 15]. Several studies reported a reduced abundance of butyrate-producing bacteria and lower fecal and luminal butyrate concentrations in IBD patients [16–19]. In addition, reduced levels of the butyrate transferase gene have been shown in IBD patients compared to healthy individuals [19, 20]. However, the majority of previous research has primarily focused on the alteration of butyrate-producing bacteria utilizing butyrate transferase as the dominant terminal enzyme for butyrate production such as *Faecalibacterium prausnitzii* in clinical studies and *in vivo* studies [16, 17, 21–24].

Interleukin-10 (IL10) is a cytokine secreted by the innate and adaptive immune cells including macrophages, dendritic cells, T cells, and B cells. IL10 plays a crucial role in maintaining tissue homeostasis through its anti-inflammatory properties, which include the modulation of innate and adaptive immunity [25, 26]. IL10 knockout mice have been used as a IBD mouse model, as they develop spontaneous chronic colitis, which takes 2-3 months [27]. The commensal gut microbiota is necessary to initiate and sustain intestinal inflammation, making it a suitable model to study changes in gut microbiota during the onset and development of IBD [27, 28]. Moreover, human infantile onset IBD can result from rare mutations in IL10 or IL10 receptor, suggesting the role of IL10 pathway in maintaining intestinal homeostasis and preventing the intestinal inflammation [29–32]. Several studies have reported dysbiosis in IBD mouse models including IL10 knockout mice [7, 33]. However, there is limited research on changes in butyrate-producing bacteria in experimental IBD mouse models at various taxonomic levels.

Here, this study aims to characterize changes in butyrate-producing bacteria at various taxonomic levels in a mouse model of inflammatory bowel disease, specifically using IL10 knockout mice. We demonstrate diminished butyrate oxidation while elevated glycolysis in 9-week IL10 knockout primary colonocytes. Furthermore, we find a significant increase in butyrate kinase and reduced levels of butyrate transferase in 9-week IL10 knockout mice. These data align with increased abundance of *Coprococcus comes* that use butyrate kinase for butyrate production and decreased abundance of *Faecalibacterium prausnitzii* that use butyrate transferase for butyrate production along with the decreased acetate levels in in 9-week IL10 knockout mice. These data suggest that changes in butyrate-producing bacteria can be observed prior to the onset of colitis as supported in our IL10 knockout model of spontaneous colitis. Furthermore, changes in bacteria that harbor butyrate kinase represent a potential compensatory mechanism to increase butyrate production through an alterative non cross-feeding pathway.

## Materials and methods

### Animals

Male wild type (WT) C57BL/6 (BL6) mice obtained from the Jackson Laboratories (Bar Harbor, ME) and IL10 knockout (IL10 KO) BL6 mice were provided by Dr. Jeremiah Gene Johnson (University of Tennessee, Knoxville). The mice were housed in a humidity-controlled room at 22-23 °C on 12-hour light/dark cycle under specific pathogen-free condition at the University of Tennessee, Knoxville. All experiments were conducted following the policies approved by the Institutional Animal Care and Use Committee at the University of Tennessee, Knoxville.

The mice in this study were euthanized at 9 weeks old and their cecum, colon, and fecal colon contents were collected for primary colonocytes isolation, short chain fatty acids analysis or bacterial DNA isolation.

### Isolation of primary colonocytes

Colonocytes were isolated as previously used to exclude cell types except for colonocytes, including immune cells, smooth muscle, and enteric neurons [11, 34, 35]. After sacrificing each mouse, the entire colon was dissected and thoroughly washed with sterile phosphate-buffered saline (PBS). The tissue was placed in the isolation buffer (1X PBS with 5 mM EDTA and 1% fetal bovine serum (FBS)) and incubated for 20 min at 37 °C with gentle agitation. After 20 minutes, colonic tissue was removed, and isolated colonocytes were collected through centrifugation for flux experiments.

### Flux Experiments

The Seahorse XFe24 Analyzer (Agilent Technologies, CA) was used for real-time measurements of oxygen consumption rate (OCR) and glycolytic proton efflux rate (glycoPER). The butyrate oxidation assay was optimized in our previous work [36]. Isolated colonocytes were seeded onto the cell culture microplate. The butyrate oxidation assay was conducted in 1X Krebs-Henseleit Buffer (KHB), including 2% FBS, 5 mM glucose and 500 μM carnitine, following the addition of 5 mM sodium butyrate (Sigma, #B5887), 50 mM 2-deoxyglucose (2DG) (Sigma, #5875), and 10% sodium azide (Sigma, S-8032).

The glycolytic rate assay was performed following the manufacturer’s instruction using XF Assay medium (Agilent, 103575-100) supplemented with 2% FBS, 2 mM L-glutamine, 10 mM glucose, and 1 mM sodium pyruvate. Sequential injections of 0.5 μM Rotenone/Antimycin and 50 mM 2DG were used during the assay. After each assay, cell lysates were prepared using 1X RIPA buffer (Cell Signaling, 9806s), and protein levels were quantified using a Pierce BCA Protein Assay kit (Thermo Fisher, #PI23228) to normalize each assay.

### Quantification of short chain fatty acids

Fecal colonic contents were collected in 1.5 ml Eppendorf tubes and frozen at -80 °C until the day of analysis to determine the concentrations of SCFAs: acetate, propionate, and butyrate. The samples were prepared as described previously with a few modifications [37]. Fecal colonic contents stored at -80 °C were thawed on wet ice, and 40 mg of samples were transferred to fresh tubes. Subsequently, 400 µl of distilled water was added to achieve a homogeneous suspension. The samples were mixed with a metal spatula, centrifuged at 13000 rpm for 5 minutes and 200 µl of the suprnatant from the homogenates were transferred into a new 1.5 ml tube. To this homogenate, 200 µl of an organic solvent mixture was added, comprising N-butanol, tetrahydrofuran and acetonitrile in a ratio of 50:30:20, along with 40 ul of 0.1 M HCl, 20 mg of citric acid, and 40 mg of sodium chloride. The samples were vigorously homogenized by vortexing for 1 minute and centrifuged at 13,000 rpm at room temperature for 10 minutes. The supernatant was pipetted and transferred to chromatographic vials.

Gas chromatography (GC) analysis was employed to quantitate acetate, propionate, and butyrate in fecal colonic contents using a GC instrument (Agilent Technologies; 6890). The GC was equipped with a DB-23 column (30 m × 0.25 mm × 0.25 μm film thickness; Agilent Technologies) and a flame ionization detector (FID). The GC-FID system parameters were as follows: split ratio, 1:10; carrier gas, helium; flow rate, 1ml/min; inlet temperature, 250 C. The oven parameters were as follows; initial temperature, 100 °C for 7 mins; 25 °C/min to 200 °C, held for 5 minutes. The FID parameters were as follows: temperature, 260 °C; hydrogen flow, 30 ml/min; synthetic air, 250 ml/min.

Data were collected with GC Chemstation Rev. A 10.02 [1757] software (Agilent Technologies). The samples’ SCFA concentrations were quantified using external calibration equations prepared with a mixture of acetate, propionate, and butyrate that were prepared according to the liquid-liquid extraction procedures used above for the fecal colonic contents of the samples.

### Bacterial DNA extraction from cecum, colon, colon contents and quantitative real-time PCR

Bacterial DNA from cecum, colon, and fecal colonic contents of WT and IL10 KO mice was isolated using DNeasy PowerSoil Pro Kit (QIAGEN Sciences Inc, Germantown, MD) following the manufacturer’s instruction. The extracted DNA was stored at -80 °C for further analysis. DNA concentration was measured using the NanoDropND-1000 spectrophotometer (NanoDrop Technologies, Wilmington, DE).

The primers used in this study are listed in Table 1. Bacterial 16S rRNA were quantified in the cecum, colon, and fecal colon contents using the Real Time PCR 7300system (ABI, Foster City, CA). The amplifications were performed in a final reaction volume of 15 ul containing 2X SYBR mix (Life technologies, #25742), each forward and reverse primer, bacterial DNA, and molecular biology grade water. The amplification protocol was done as follows; one cycle at 95°C for 2 min, followed by 35 cycles of 95°C for 45 seconds, 54°C for 45 seconds, and 72°C for 45 seconds. Fluorescent products were measured at the last step of each cycle.

**Table 1.**
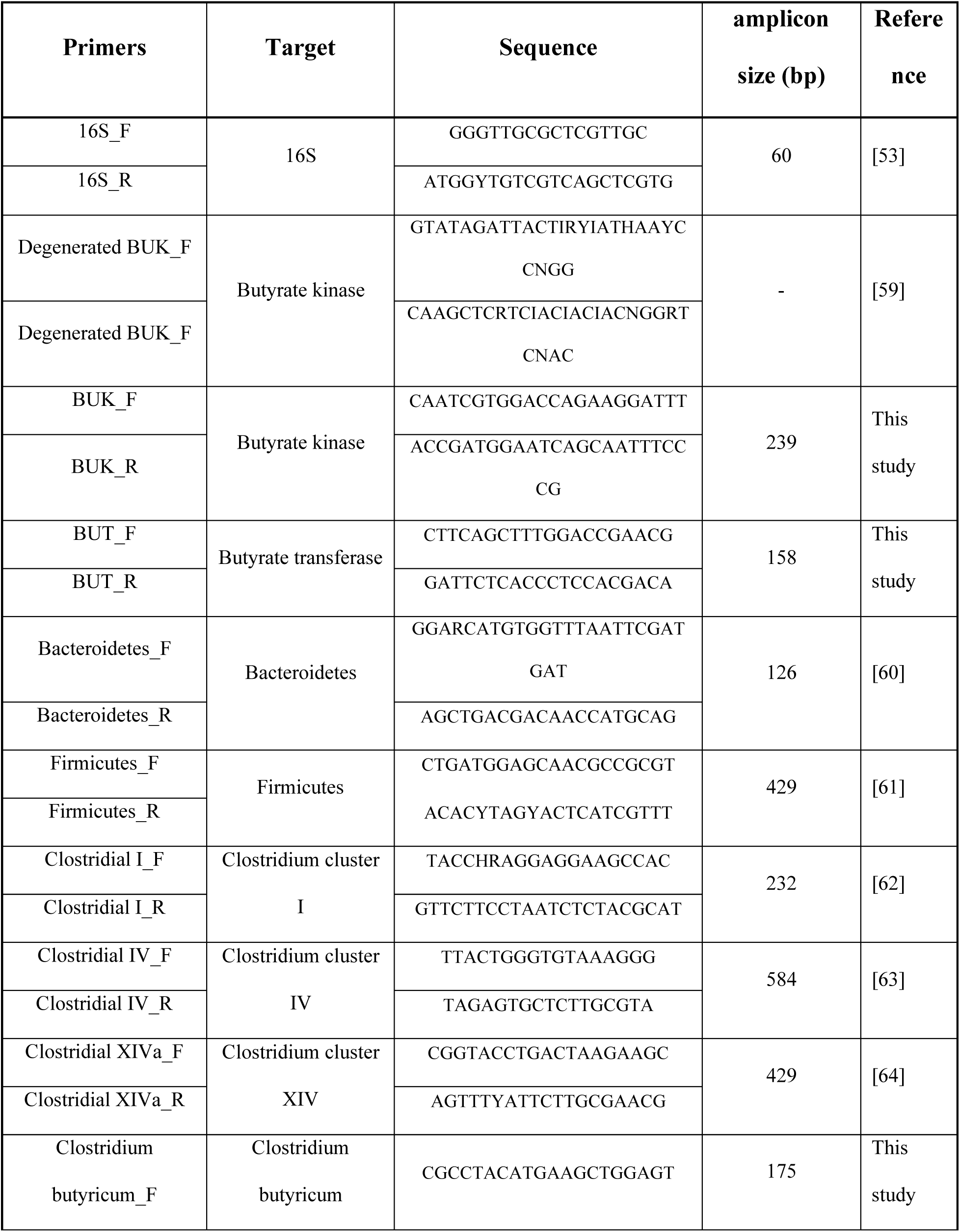
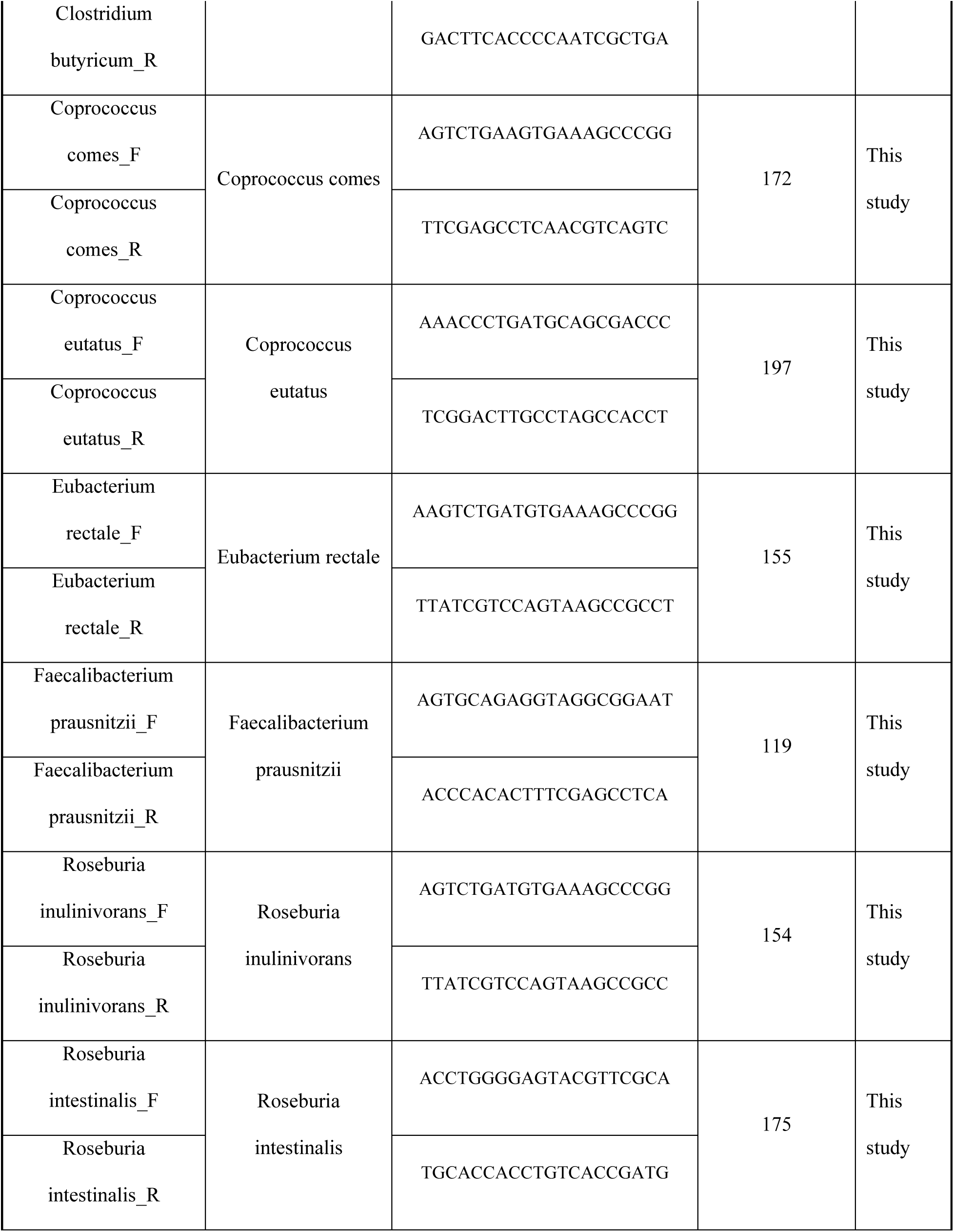
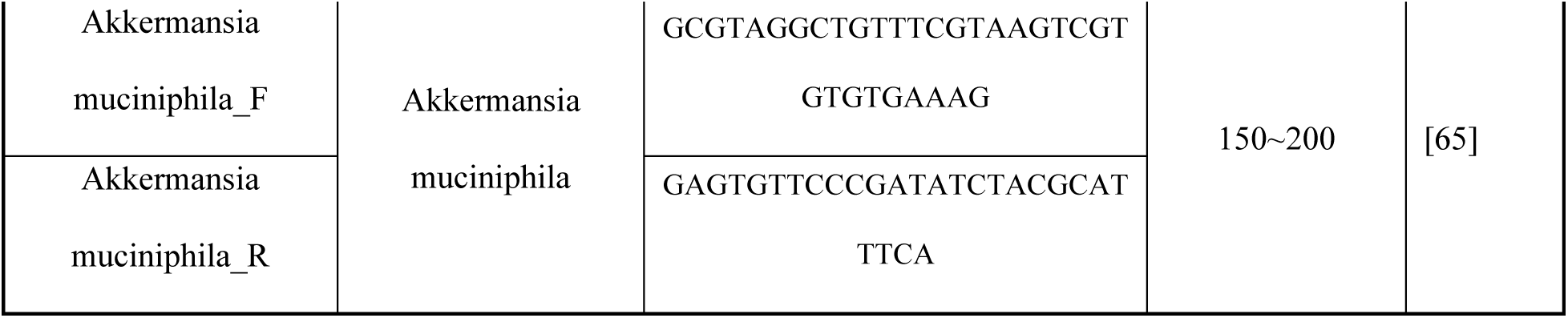
Primers used for qPCR analysis by 16S rRNA.

### Statistical Analysis

Statistical analysis was conducted using GraphPad Prism 10. t-test or ANOVA was used to test for differences between experimental groups, followed by a Tukey post-hoc test for the ANOVA. All data are expressed as mean ± SEM. Significant differences are indicated.

## Results

### Signs of Increased Inflammation at 9-weeks in IL10 Knockout Mice

Previous work has shown that IL10 KO mice develop spontaneous colitis starting at 10-12 weeks of age [27]. The animal facility and gastrointestinal microbiome of the mice are big factors that influence the timing of this event [38]. To test whether there are significant biometric indicators that might indicate the development of colitis, body weight, spleen and cecum weight, and colon length were measured in WT and 9-week-old pre-colitis IL10 KO mice. There was not a significant difference between body weight in IL10 KO mice at 9 weeks of age as compared to WT mice (**Figure 1A**). However, both spleen (**Figure 1B**) and cecum (**Figure 1C**) weights were higher in IL10 KO mice, with the increased spleen weight representing an inflammatory metric. Colon length was significantly decreased at 9 weeks in IL10 KO mice compared to WT mice (**Figure 1D**). Since IL10 KO mice showed no difference in body weight and changes in other parameters suggests that these mice are in a heightened inflammatory state but haven’t developed colitis.

**Figure 1.**
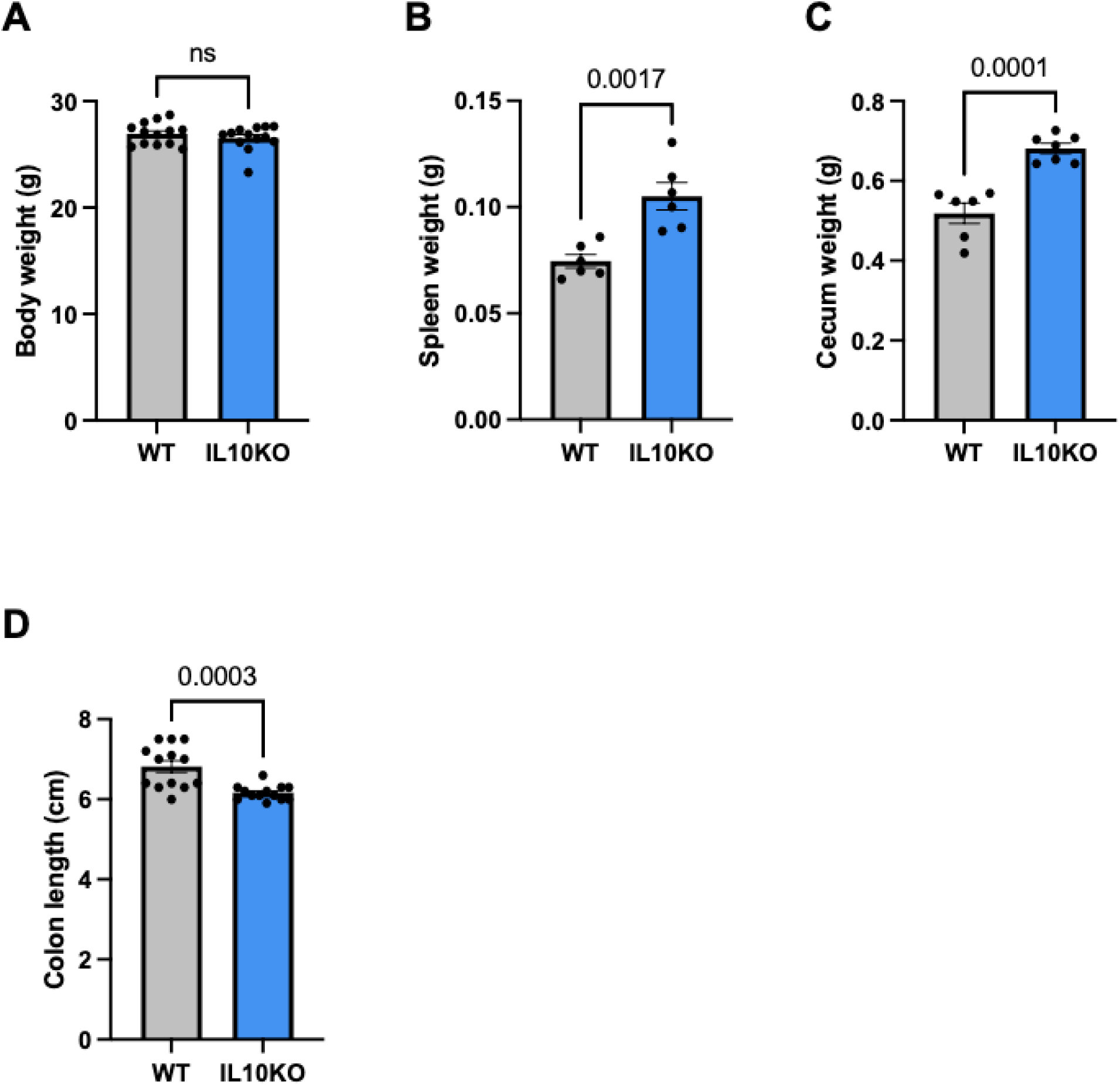
Biometrics in IL10 Knockout Mice. **(A)** Body weight, **(B)** spleen weight, **(C)** cecum weight, **(D)** colon length of wildtype (WT) mice and IL10 knockout (IL10 KO) mice. Error bars indicate the mean ± SD (n = 6 or 13). Significant differences are shown.

### Diminished Response to Butyrate in IL10 Knockout colonocytes

Colitis has been associated with a decrease in the oxidation of the short chain fatty acid butyrate [39–41]. Based on the fact that IL10 KO mice develop spontaneous colitis, we postulated whether colonocytes isolated prior to the development of colitis would also show reduced butyrate oxidation. Colonocytes isolated from 9-week-old IL10 KO mice showed a decrease in the oxidation of butyrate as compared to the WT BL6 counterpart colonocytes (**Figure 2A and B**). In contrast, IL10 KO colonocytes had a higher glycolytic rate as judged by the elevated glycoPER or protons derived from glycolysis as opposed to carbon dioxide (**Figure 2C**). This was associated with an increase in the basal glycolysis (**Figure 2D**) and compensatory glycolysis (**Figure 2E**). These findings align with a decreased butyrate oxidation **(Figure S1A)** and an increased basal glycolysis **(Figure S1B)** and compensatory glycolysis **(Figure S1C)** in 5-week IL10 KO colonocytes. Taken together, these data suggest that IL10 KO colonocytes reduce butyrate oxidation at the expense of increasing glycolysis.

**Figure 2.**
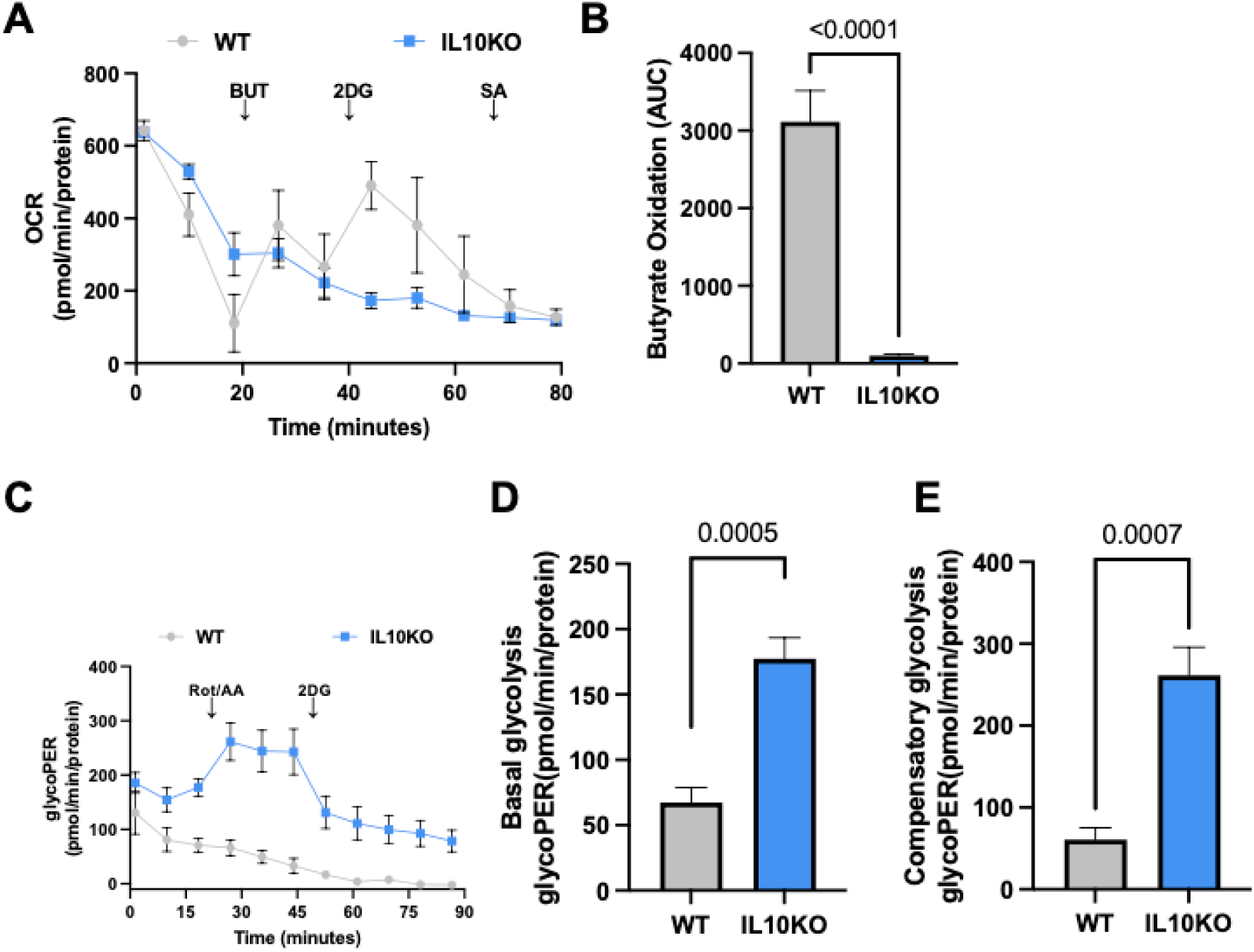
IL10 Knockout Colonocytes Diminished Response to Butyrate and Increased Response to Glucose. **(A)** Oxygen consumption rate (OCR) in wildtype (WT) mice and IL10 knockout (IL10 KO) mice with butyrate (5 mM). Total contribution of butyrate toward OCR is determined after injection of 2-deoxyglucose (2DG). Butyrate oxidation **(B)** represents the area under the curve (AUC) from OCR measurements taken after 2DG injection but before sodium azide (SA) injection. Data points represent the OCR per condition for butyrate oxidation measurements. Contribution of glycolysis to proton efflux rate (glycoPER) **(C)**, basal glycolysis **(D)**, and compensatory glycolysis **(E)** in WT mice and IL10 KO mice. Data points represent the glycoPER per condition for glycolysis measurements. Error bars indicate the mean ± SD (n = 4 or 5). Significant differences are shown.

### Decreased Colonic Acetate Levels in IL10 Knockout Mice

To test whether these pre-colitis mice showed any decrease in the major short chain fatty acids, acetate, propionate, or butyrate, we collected fecal colonic contents from 9-week-old IL10 KO or WT BL6 mice and measured SCFAs using gas chromatography. Interestingly, we found that only acetate was significantly decreased in IL10 KO mice (**Figure 3A**). The concentration of propionate (**Figure 3B**) or butyrate (**Figure 3C**) was not different between the groups. The fact that butyrate was not also diminished in IL10 KO was surprising and we next sought to determine if butyrate-producing bacteria were altered in the gut microbiome of IL10 KO mice.

**Figure 3.**
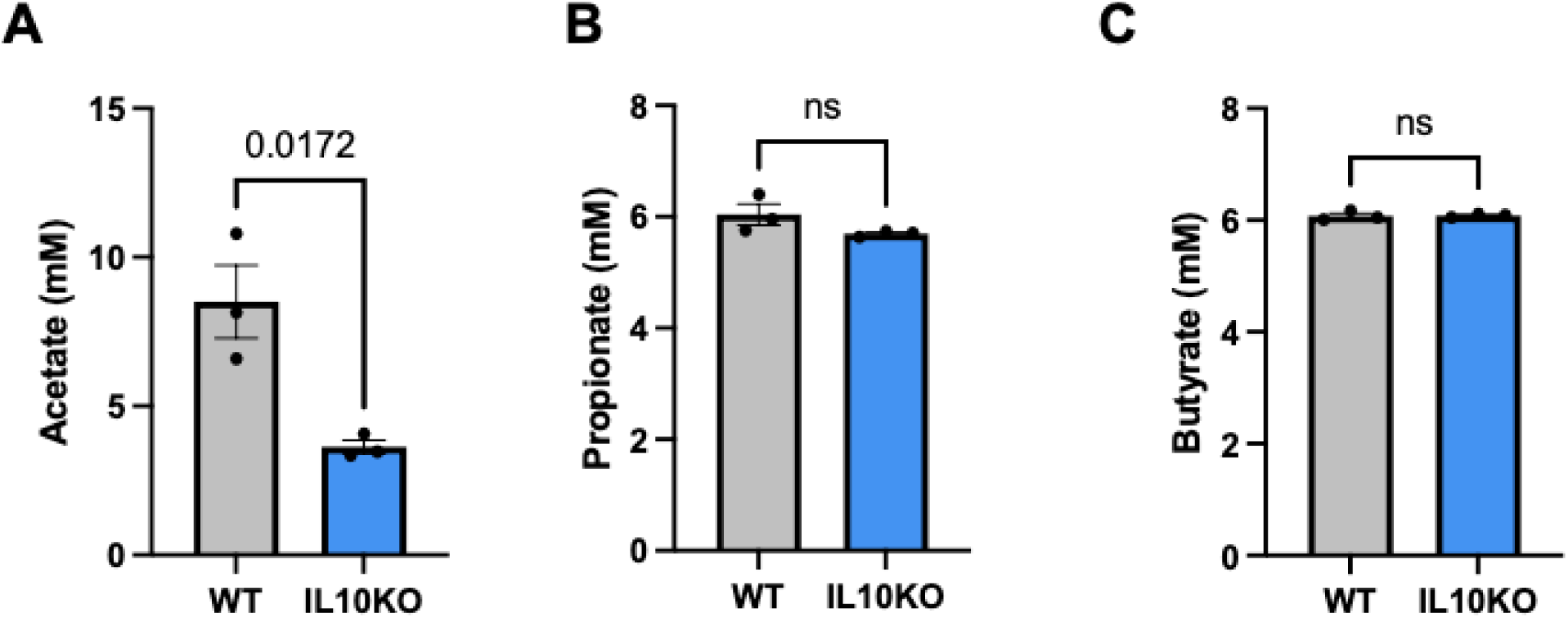
Profile of Short Chain Fatty Acids in IL10 Knockout Mice. (**A**) Acetate, (B) propionate, and (C) butyrate levels in the fecal colonic contents of wildtype (WT) mice and IL10 knockout (IL10 KO) mice. Error bars indicate the mean ± SD (n = 3). Significant differences are shown.

### Alteration in the Levels of Terminal Enzymes for Butyrate Production in IL10 Knockout Mice

Butyrate kinase (Buk) and butyryl CoA:acetate CoA-transferase (But), also called butyrate transferase, are two important enzymes in the bacterial butyrate synthesis pathway. Butyrate kinase catalyzes the butyryl phosphate to butyrate and butyrate transferase utilizes acetate to accept the CoA from butyryl-CoA to form butyrate, which also represents a cross-feeding pathway meaning the presence of acetate is crucial in its formation **(Figure 4A**). Unexpectedly, when butyrate kinase levels (representative of the amount of butyrate-producing bacteria that harbor this enzyme) relative to total 16s was measured in gut microbiome, including the cecum (**Figure 4B**), colon (**Figure 4C**), and fecal colonic contents collected in the colonic lumen (**Figure 4D**), 9-week IL10 KO mice showed significantly higher Buk compared to age matched WT BL6 controls. Comparatively, 5-week old IL10 KO mice did not have a difference in Buk compared to control **(Figure S1D)**. On the other hand, butyrate transferase levels in 9-week-old IL10 KO mice were significantly diminished in cecum (**Figure 4E**), and fecal contents (**Figure 4F**), while trending toward reduced levels in colon (**Figure 4G**). This increase in Buk and decrease in But in pre-colitis mice may be associated with the reduced acetate levels, which could induce a compensatory increase in the alternative butyrate kinase pathway.

**Figure 4.**
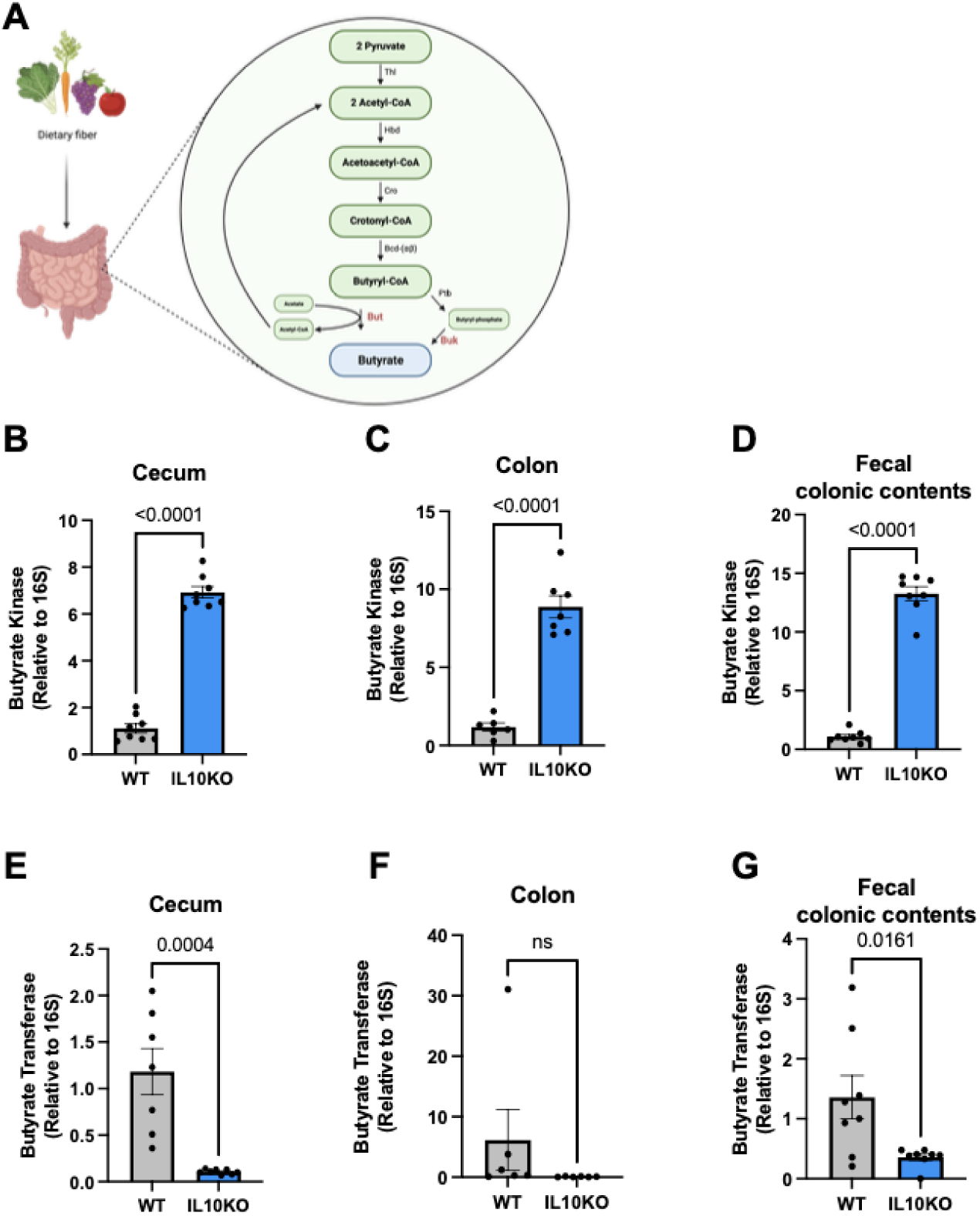
Altered Levels of the Terminal Enzymes for Butyrate Production in IL10 Knockout Mice. Metabolic pathway for butyrate production **(A).** Thl; thiolase; Hbd, 2-hydroxybutyryl-CoA dehydrogenase; Cro, crotonase; Bcd-(αβ). Butyryl-CoA dehydrogenase (including electron transfer protein αβ subunit); Buk; butyrate kinase; But, butyryl-CoA:acetate CoA transferase. Abundance of buk was measured in the cecum **(B)**, colon **(C)**, and fecal colonic contents **(D)** of wildtype (WT) mice and IL10 knockout (IL10 KO) mice. Abundance of but was measured in the cecum **(E)**, colon **(F)**, and fecal colonic contents **(G)** of WT mice and IL10 KO mice. Error bars indicate the mean ± SD (n = 6 or 8). Significant differences are shown.

### Diminished Butyrate-producing Bacteria at the Phylum and Genus Level in IL10 Knockout Mice

Changes in butyrate-producing bacteria has been reported in the profile of gut microbiota in colitis [20, 42, 43]. To investigate whether there is a significant change in butyrate-producing bacteria at the phylum and genus level in IL10 KO mice, we measured the phyla Bacteroidetes and Firmicutes and the genera *Clostridial I*, *Clostridial IV*, and *Clostridial XIVa* in DNA samples collected from the cecum, colon, and fecal colonic contents. At the phylum level, the abundance of Firmicutes (representative of most of butyrate-producing bacteria) was significantly decreased in the cecum, colon, and fecal colonic contents of IL10 KO mice compared to that in WT BL6 controls (**Figure 5B**). However, the levels of Bacteroidetes (representative of non-butyrate-producing bacteria) were significantly increased in the cecum and fecal colonic contents with a trend towards increased levels in the colon (**Figure 5A**). At the genus level, the abundance of *Clostridial I* and *IV*, well-established butyrate-producing bacteria populations, was significantly lower in IL10 KO mice than in WT BL6 mice (**Figure 5C, 5D**) with a trend toward decreased levels of *Clostridial IV* in the colon of IL10 KO mice. However, there was no significant difference in the abundance of *Clostridial XIVa*, another well-established butyrate-producing bacteria population, in IL10 KO mice, except for lower levels of *Clostridial XIVa* in the cecum of IL10 KO mice (**Figure 5E**).

**Figure 5.**
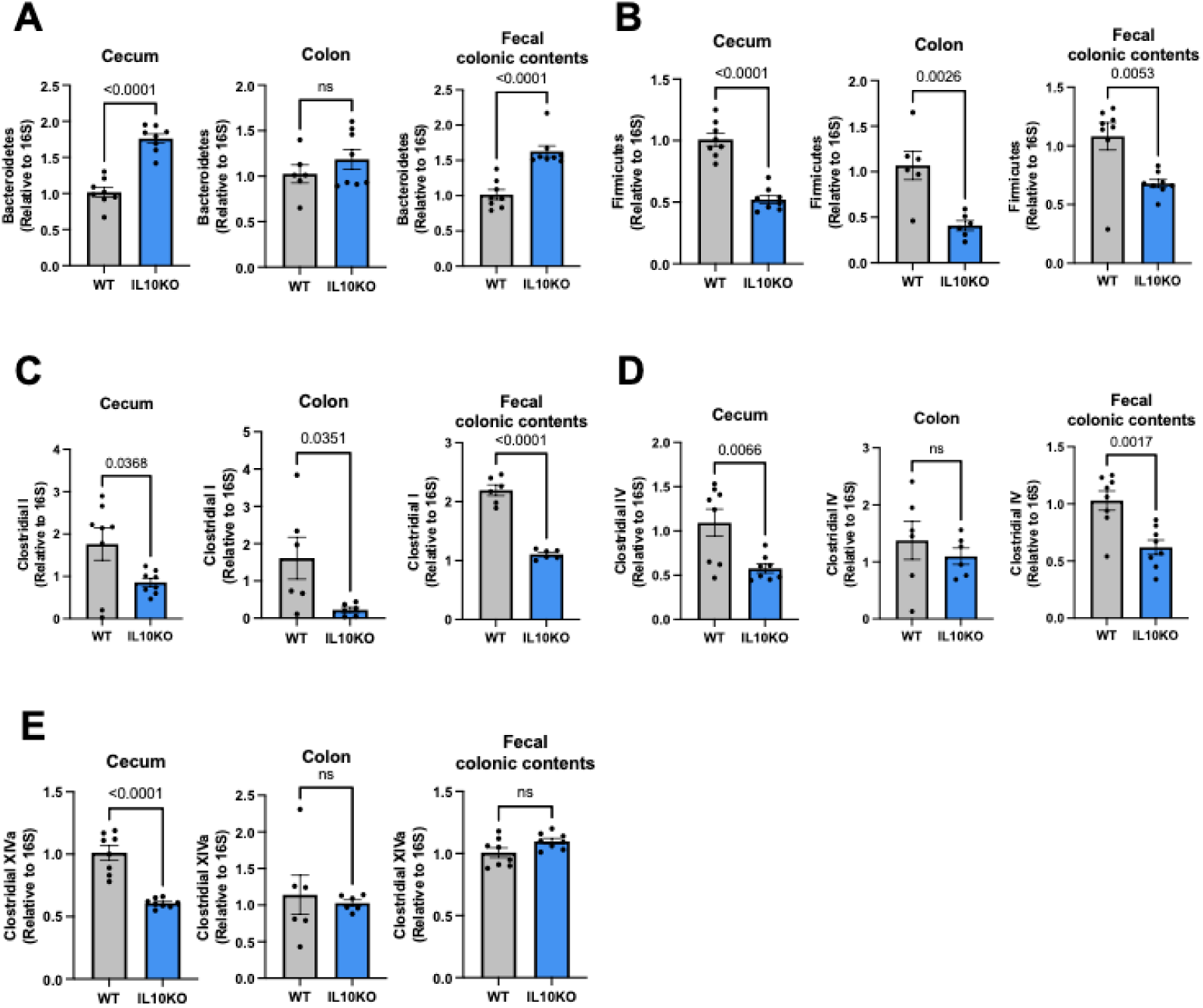
Changes in Butyrate-Producing Bacteria at the Phylum and Genus level in IL10 Knockout Mice. **(A)** Abundance of the phylum bacteroidetes was measured in the cecum, colon, and fecal colonic contents of wildtype (WT) mice and IL10 knockout (IL10 KO) mice. **(B)** Abundance of the phylum firmicutes was measured in the cecum, colon, and fecal colonic contents of WT mice and IL10 KO mice. **(C)** Abundance of the genus Clostridial I was measured in the cecum, colon, and fecal colonic contents of WT mice and IL10 KO mice. **(D)** Abundance of the genus Clostridial IV was measured in the cecum, colon, and fecal colonic contents of WT mice and IL10 KO mice. **(E)** Abundance of the genus Clostridial XIV was measured in the cecum, colon, and fecal colonic contents of WT mice and IL10 KO mice. Error bars indicate the mean ± SD (n = 6 or 8). Significant differences are shown.

### Changes of Butyrate-Producing Bacteria Harboring Butyrate Kinase in IL10 Kncokout Mice

A majority of studies have focused on using butyrate transferase as a biomarker to investigate the butyrate-producing capacity, since but-harboring bacteria were more dominant than buk-harboring bacteria in humans [44]. However, a recent study observed that Buk seems to be more dominant for butyrate synthesis in the mouse host, although both buk and but are prevalent in the mouse microbiome for butyrate production [45]. Additionally, we found unexpected increase of butyrate kinase in IL10 KO mice (**Figure 3A, 3B, 3C**) and postulated whether there is a significant increase in butyrate-producing bacteria that encode butyrate kinase at the species level in IL10 KO mice. Interestingly, the amounts of *Coprococcus comes*, which belongs to *Clostridial XIVa*, in the cecum, colon, and fecal colonic contents of IL10 KO mice, were significantly higher than that in WT BL6 mice (**Figure 6B**). This trend is consistent with the levels of butyrate kinase in IL10 KO mice (**Figure 4B, 4C, 4D**). However, the abundance of *Clostridium butyricum*, one of *Clostridial I* members, in the cecum and colon of IL10 KO mice was significantly lower than that in WT mice, with a trend towards decreased levels in the fecal colonic contents (**Figure 6A**). This species’ abundance in IL10 KO mice was matched to the abundance of the genus to which it belongs (**Figure 5C**).

**Figure 6.**
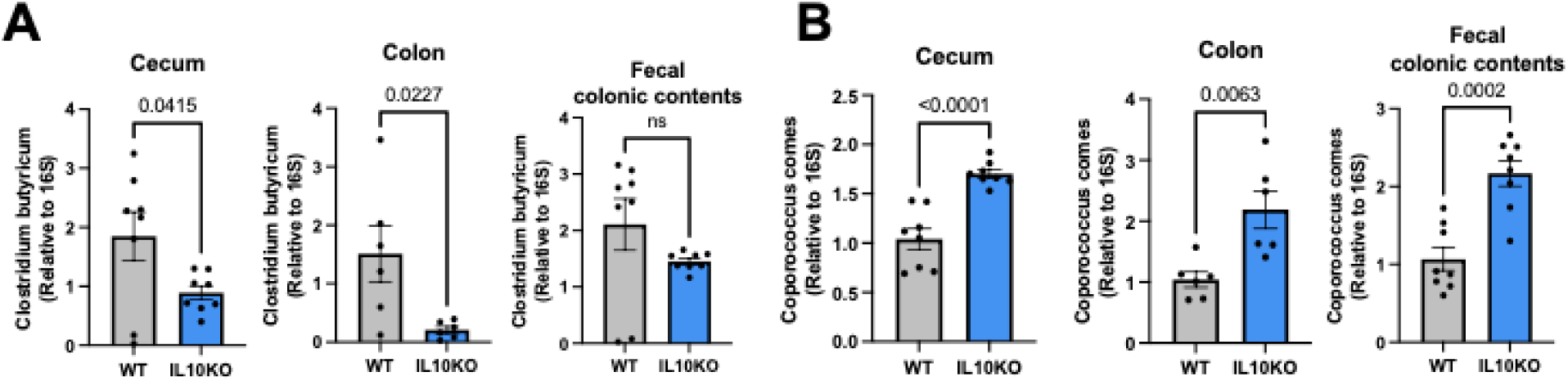
Altered Abundance of Butyrate-Producing Harboring Butyrate Kinase at the Species level in IL10 Knockout Mice. **(A)** Abundance of the species Clostridium butyricum was measured in the cecum, colon, and fecal colonic contents of wildtype (WT) mice and IL10 knockout (IL10 KO) mice. **(B)** Abundance of the species Coprococcus comes was measured in the cecum, colon, and fecal colonic contents of WT mice and IL10 KO mice. Error bars indicate the mean ± SD (n = 6 or 8). Significant differences are shown.

### Altered Butyrate-producing Bacteria that Harbor Butyrate Transferase in IL10 Knockout Mice

The study next delved into the abundance of several major butyrate-producing Bacteria at the species level that encode butyrate transferase, including *Coprococcus eutatus*, *Eubacterium rectale*, *Faecalibacterium prauscnitzii*, *Roseburia inulivorans*, and *Roseburia intestinalis*. In the cecum, colon, and fecal colonic contents in IL10 KO mice, the amounts of *Faecalibacterium prauscnitzii* were significantly lower than those in IL10 KO mice **(Figure 7B)**. This pattern aligns with the amounts of *Clostridial IV*, to which *Faecalibacterium prauscnitzii* belongs, and the levels of butyrate transferase, in IL10 KO mice **(Figure 3D, 3E, 3F, 5D)**. Additionally, *Coprococcus eutatus*, exhibiting *Buk* and *But*, showed lower levels in the cecum, colon, and fecal colonic contents of IL10 KO mice compared to WT BL6 mice **(Figure S2B)**. However, there were no significant differences of *Roseburia inulinvorans* and *Roseburia intestinalis* in the cecum, colon, and fecal colonic contents between WT BL6 mice and IL10 KO mice **(Figure 7C, 7D)**. Unexpectedly, the abundance of *Eubacterium rectale* was increased in IL10 KO mice compared to WT mice, except for no alteration in the colon of IL10 KO mice **(Figure 7A)**. The differing abundance trend of *Coproccous eutatus*, *Eubacterium rectale*, *Roseburia inulinvorans*, and *Roseburia intestinalis*, which belong to *Clostridial XIVa*, may be associated with the lack of significant differences between WT mice and IL10 KO mice in the abundance of *Clostridial XIVa* in the colon and fecal colonic contents.

**Figure 7.**
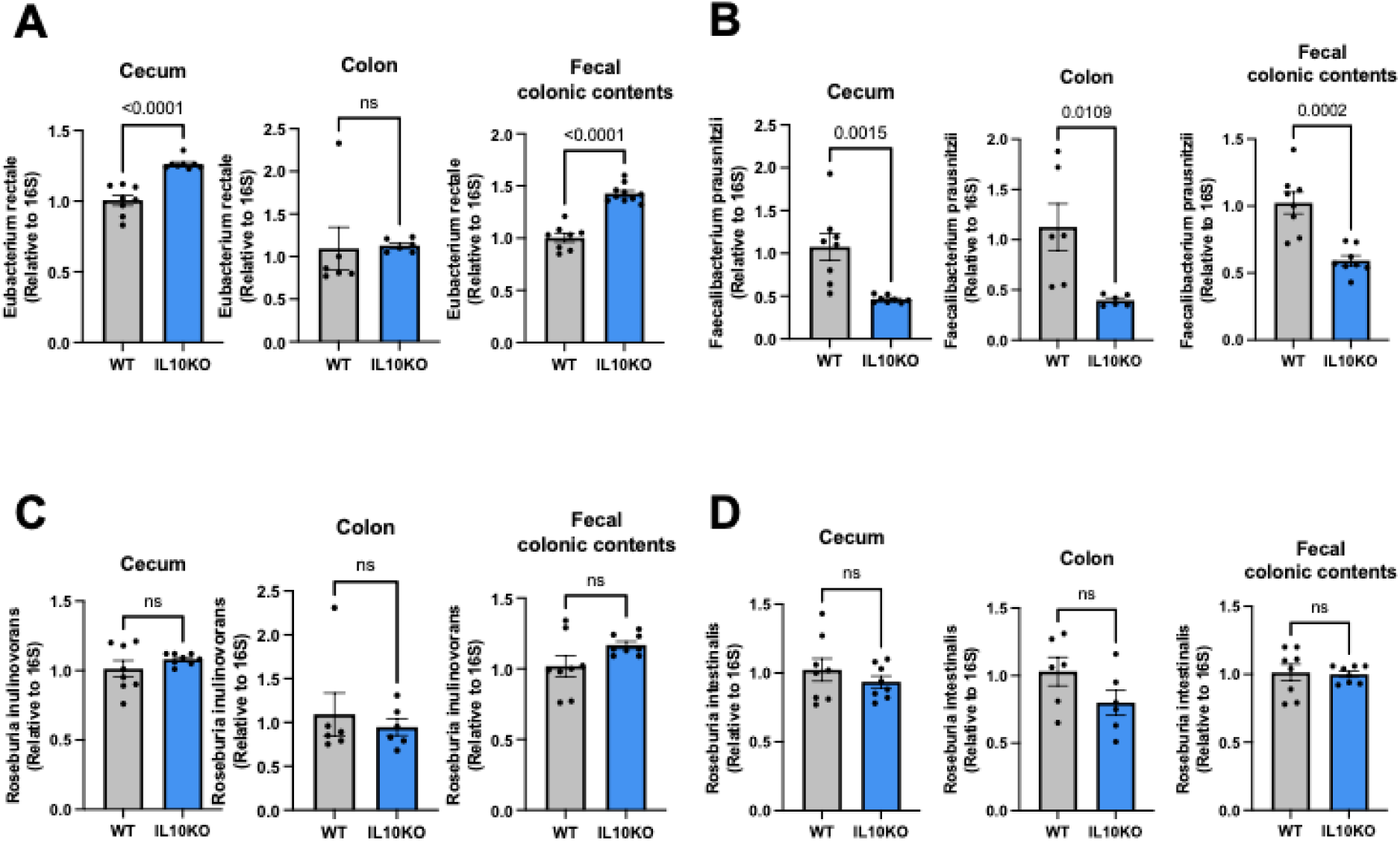
Abundance of Butyrate-Producing Harboring Butyrate Transferase at the Species level in IL10 Knockout Mice. **(A)** Abundance of the species Eubacterium rectale was measured in the cecum, colon, and fecal colonic contents of wildtype (WT) mice and IL10 knockout (IL10 KO) mice. **(B)** Abundance of the species Faecalibacterium prauscnitzii was measured in the cecum, colon, and fecal colonic contents of WT mice and IL10 KO mice. **(C)** Abundance of the species Roseburia inulinvorans was measured in the cecum, colon, and fecal colonic contents of WT mice and IL10 KO mice. **(D)** Abundance of the species Roseburia intestinalis was measured in the cecum, colon, and fecal colonic contents of WT mice and IL10 KO mice. Error bars indicate the mean ± SD (n = 6 or 8). Significant differences are shown.

## Discussion

This study aims to characterize changes in butyrate-producing bacteria at various taxonomic levels in a pre-colitis mouse model, specifically using IL10 KO mice. Our findings reveal diminished butyrate oxidation and elevated glycolysis in 9-week IL10 KO primary colonocytes. These findings align with previous research reporting impaired butyrate oxidation and enhanced glycolysis in the colonic mucosa of patients with UC [39–41]. Additionally, one study has reported the decreased expression of genes related to the butyrate oxidation in CD colonic mucosal biopsies [46]. This metabolic shift may be attributed to a compensatory mechanism resulting from the energy deficiency hypothesis, potentially linked to the activation of AMPK in IL10 KO mice [41, 47]. Thus, the reduced capacity of colonocytes to oxidize butyrate may be related to a critical metabolic deficit observed in UC patients.

Furthermore, we observe a significant reduction in BPB at the phylum Firmicutes, and identified decreased some BPB species including *Clostridium butyricum*, *Coprococcus eutatus*, *Faecalibacterium prausnitzii*, with decreased levels of the terminal enzyme for butyrate production, butyrate transferase. However, other BPB species, including *Coprococcus comes*, Eubacterium rectale and the alternative terminal enzyme for butyrate production, butyrate kinase, showed higher levels in IL10 KO mice.

Butyrate-producing bacteria belongs to the phylum Firmicutes, the families *Lachnospiraceae* and *Ruminococcaceae*, and the genus *clostridial I*, *IV* and *XIVa*. Healthy individuals have shown higher abundance of butyrate-producing bacteria compared to IBD patients. However, an imbalance in gut microbiota, specifically butyrate-producing bacteria, is commonly observed in individuals with IBD patients and experimental models [17, 18]. In our study, IL10 KO mice showed lower levels of Firmicutes and *Faecalibacterium prausnitzii* and higher levels of Bacteroidetes, as previously reported in IBD patients and experimental colitis model [21, 48–51]. This may be due to the IL10 deficiency and lack of environmental factors that contribute to IBD. Notably, we observed higher abundance of *Coprococcus comes* which uses butyrate kinase for butyrate production in the cecum, colon, and fecal colonic contents of IL10 KO mice compared to WT BL6 mice. According to a Blast search using the sequence of the buk primer used in this study, the sequence shares 71% identity with *Coprococcus comes*, which can be linked to higher levels of butyrate kinase in IL10 KO mice **(Table 2)**. This may suggest that *Coprococcus comes* is is a major contributor to the increased butyrate kinase in IL10 KO mice.

**Table 2.**
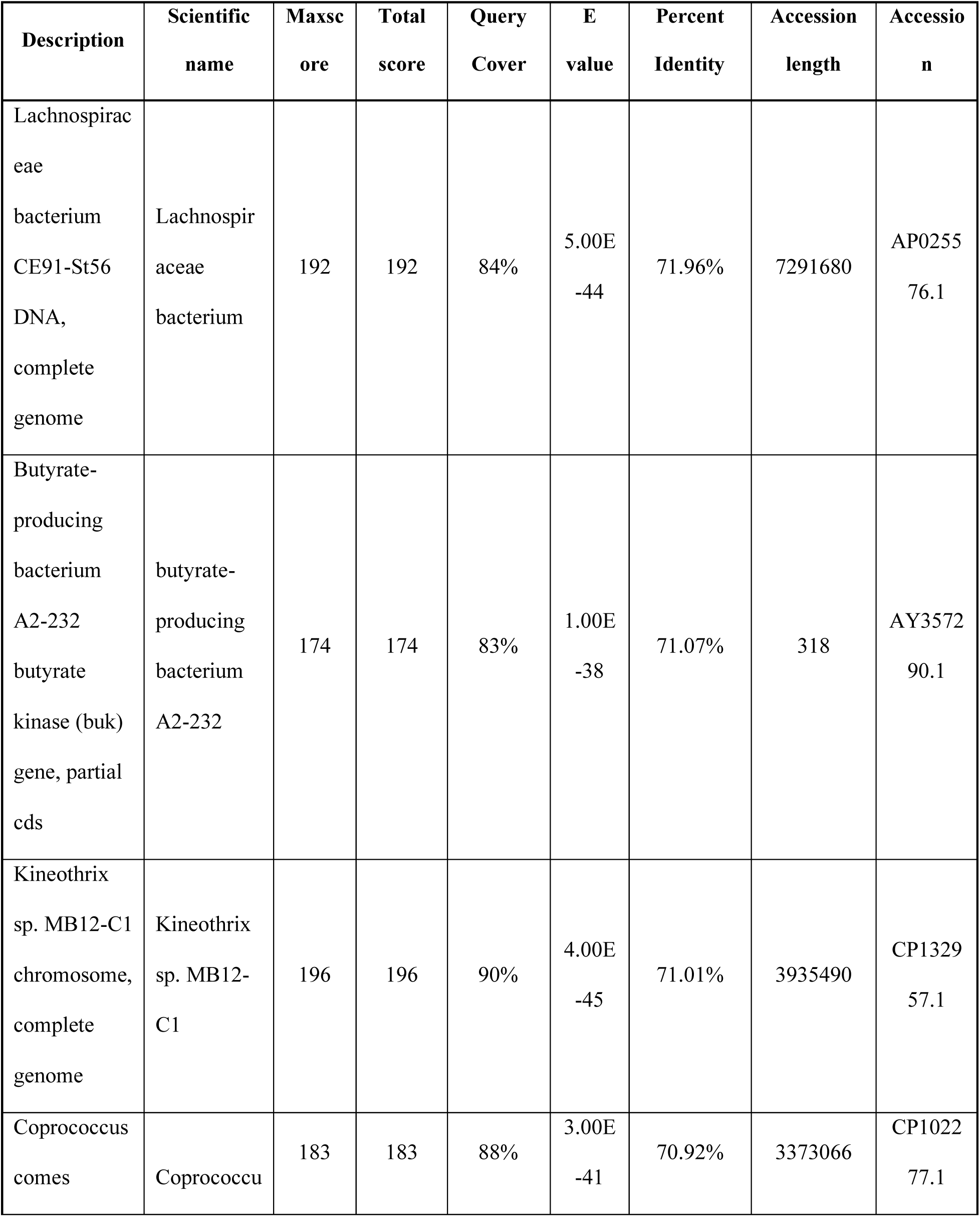
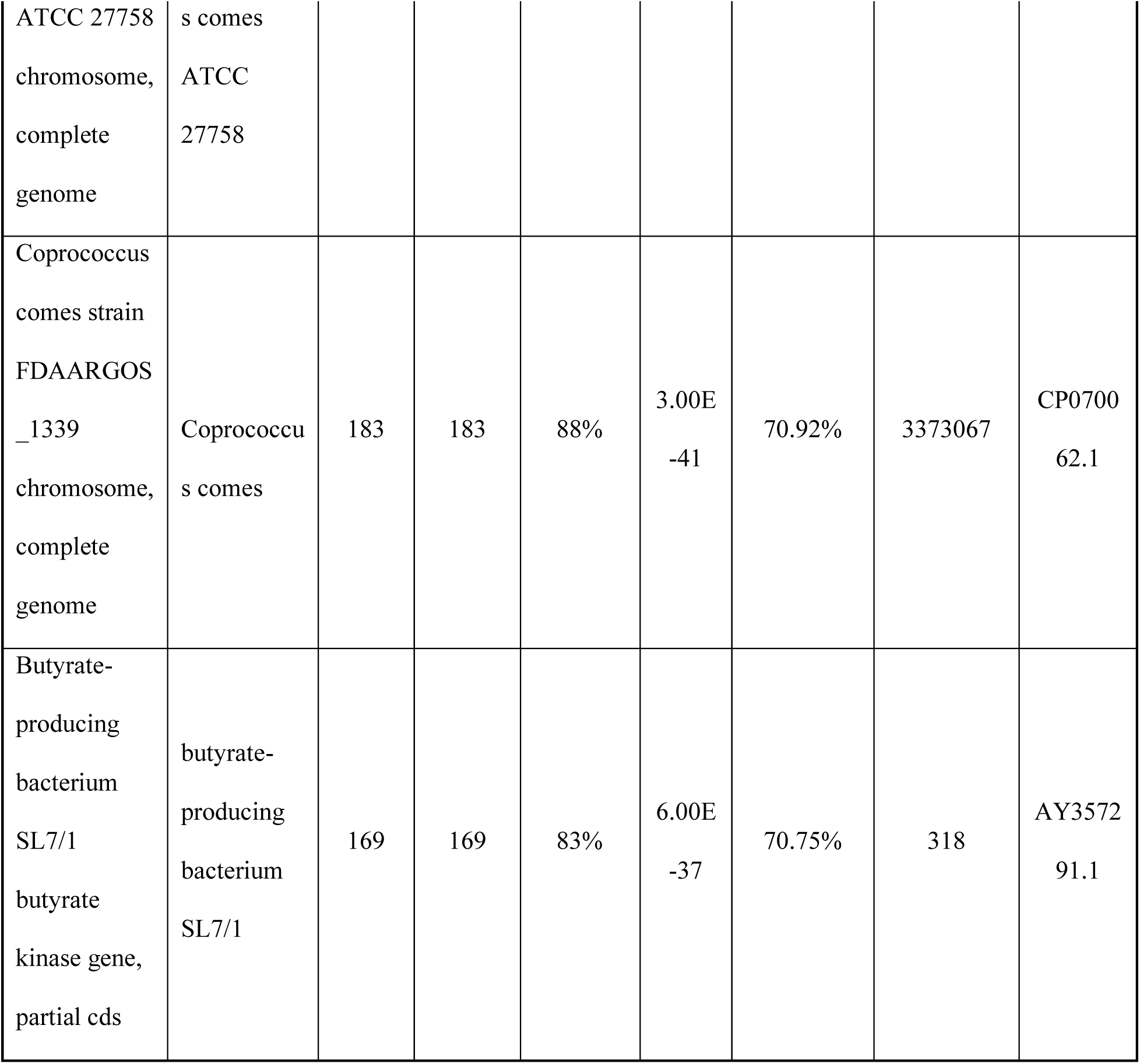
Result of Blastn search using the sequence of amplifed DNA fragment generated from the primer butyrate kinase.

Butyrate kinase and butyrate transferase are two genes that are commonly employed as biomarkers for identifying the relative levels of butyrate-producing bacteria [52, 53]. Consistent with previous data, we found lower expression butyrate transferase in IL10 KO mice [19, 20, 54]. Unexpectedly, we observed significant increase in butyrate kinase in IL10 KO mice. This result might be associated with the reduced concentrations of acetate in IL10 KO mice. The reduction in acetate levels may contribute to reduced levels of butyrate transferase, as it uses acetate as a co-substrate to produce butyrate. Furthermore, this alteration might trigger a compensatory response that increases the level of bacteria that harbor butyrate kinase to make up for the reduced butyrate transferase. Thus, this possible mechanism could explain the insignificant difference of butyrate levels between WT mice and IL10 KO mic**e**.

Another possible mechanism that explains the reduced levels of butyrate transferase and acetate in IL10 KO mice may be associated with the decreased abundance of *Akkermansia muciniphila* and *Faecalibacterium prausnitzii* in IL10 KO mice **(Figure 7B, S2A)**. *Akkermansia muciniphila* is a significant player of the colonic mucus-associated microbiota, residing within the mucus layer of the intestinal tract and liberates oligosaccharides and acetate through mucus degradation and fermentation [55, 56]. Reduced levels of this species have been observed in several gastrointestinal diseases including IBD and appendicitis [57, 58]. As such, acetate derived from *Akkermansia muciniphila* may be used as a co-substrate for butyrate production by *Faecalibacterium prausnitzii* [55]. This suggest that the decreased *Akkermansia muciniphila* could lead to decreased acetate, resulting in the reduced abundance of *Faecalibacterium prausnitzii* in IL10 KO mice.

## Conclusions

Our data suggest that two terminal enzymes involved in butyrate production, as well as the butyrate producing bacteria harboring those enzymes, are differently regulated in 9-week-old pre-colitis IL10 KO mice compared to WT mice. This study sheds lights on the potential of analyzing the composition of butyrate-producing bacteria at an early stage of IBD for diagnostic purposes.

## Acknowledgements

We would like to thank Dr. Jeremiah Johnson for providing IL10 KO BL6 mice.

## Authors’ contributions

Ji Yeon Kim helped design the experiments, performed the experiments, analyzed the data, and wrote part of the manuscript. Bohye Park, Olivia Riffey, Madison Bunch helped perform the experiments and analyzed the data. Jeremiah Johnson helped designed the experiments and analyzed the data. Zachary Burcham performed the metagenomics analysis. Dallas Donohoe designed the experiments, analyzed the data, wrote part of the manuscript, and oversaw the completion of the study.

## Availability of data and material

All generated raw data and/or analyzed data from the current study are available from the corresponding author upon reasonable request.

## Declarations

### Ethics approval and consent to participate

All experiments in this study were approved by the Institutional Animal Care and Use Committee at University of Tennesse, Knoxville.

### Consent for publication

Not applicable.

## REFERENCES

1. Kaser A, Zeissig S, Blumberg RS. Inflammatory bowel disease. Annu Rev Immunol. 2010;28:573–621; doi: 10.1146/annurev-immunol-030409-101225.

2. Parent K, Mitchell P. Cell wall-defective variants of pseudomonas-like (group Va) bacteria in Crohn’s disease. Gastroenterology. 1978;75(3):368–72.

3. Neish AS. Microbes in gastrointestinal health and disease. Gastroenterology. 2009;136(1):65–80; doi: 10.1053/j.gastro.2008.10.080.

4. Hoentjen F, Harmsen HJ, Braat H, Torrice CD, Mann BA, Sartor RB, et al. Antibiotics with a selective aerobic or anaerobic spectrum have different therapeutic activities in various regions of the colon in interleukin 10 gene deficient mice. Gut. 2003;52(12):1721–7; doi: 10.1136/gut.52.12.1721.

5. Tamagawa H, Hiroi T, Mizushima T, Ito T, Matsuda H, Kiyono H. Therapeutic effects of roxithromycin in interleukin-10-deficient colitis. Inflamm Bowel Dis. 2007;13(5):547–56; doi: 10.1002/ibd.20093.

6. Matsuoka K, Kanai T. The gut microbiota and inflammatory bowel disease. Semin Immunopathol. 2015;37(1):47–55; doi: 10.1007/s00281-014-0454-4.

7. Maharshak N, Packey CD, Ellermann M, Manick S, Siddle JP, Huh EY, et al. Altered enteric microbiota ecology in interleukin 10-deficient mice during development and progression of intestinal inflammation. Gut Microbes. 2013;4(4):316–24; doi: 10.4161/gmic.25486.

8. Morgan XC, Tickle TL, Sokol H, Gevers D, Devaney KL, Ward DV, et al. Dysfunction of the intestinal microbiome in inflammatory bowel disease and treatment. Genome Biol. 2012;13(9):R79; doi: 10.1186/gb-2012-13-9-r79.

9. Vester-Andersen MK, Mirsepasi-Lauridsen HC, Prosberg MV, Mortensen CO, Trager C, Skovsen K, et al. Increased abundance of proteobacteria in aggressive Crohn’s disease seven years after diagnosis. Sci Rep. 2019;9(1):13473; doi: 10.1038/s41598-019-49833-3.

10. Gophna U, Sommerfeld K, Gophna S, Doolittle WF, Veldhuyzen van Zanten SJ. Differences between tissue-associated intestinal microfloras of patients with Crohn’s disease and ulcerative colitis. J Clin Microbiol. 2006;44(11):4136–41; doi: 10.1128/JCM.01004-06.

11. Donohoe DR, Garge N, Zhang X, Sun W, O’Connell TM, Bunger MK, et al. The microbiome and butyrate regulate energy metabolism and autophagy in the mammalian colon. Cell metabolism. 2011;13(5):517–26.

12. Zheng L, Kelly CJ, Battista KD, Schaefer R, Lanis JM, Alexeev EE, et al. Microbial-Derived Butyrate Promotes Epithelial Barrier Function through IL-10 Receptor-Dependent Repression of Claudin-2. J Immunol. 2017;199(8):2976–84; doi: 10.4049/jimmunol.1700105.

13. Chen G, Ran X, Li B, Li Y, He D, Huang B, et al. Sodium Butyrate Inhibits Inflammation and Maintains Epithelium Barrier Integrity in a TNBS-induced Inflammatory Bowel Disease Mice Model. EBioMedicine. 2018;30:317–25; doi: 10.1016/j.ebiom.2018.03.030.

14. Falony G, Lazidou K, Verschaeren A, Weckx S, Maes D, De Vuyst L. In vitro kinetic analysis of fermentation of prebiotic inulin-type fructans by Bifidobacterium species reveals four different phenotypes. Appl Environ Microbiol. 2009;75(2):454–61; doi: 10.1128/AEM.01488-08.

15. Louis P, Flint HJ. Diversity, metabolism and microbial ecology of butyrate-producing bacteria from the human large intestine. FEMS Microbiol Lett. 2009;294(1):1–8; doi: 10.1111/j.1574-6968.2009.01514.x.

16. Eeckhaut V, Machiels K, Perrier C, Romero C, Maes S, Flahou B, et al. Butyricicoccus pullicaecorum in inflammatory bowel disease. Gut. 2013;62(12):1745–52; doi: 10.1136/gutjnl-2012-303611.

17. Machiels K, Joossens M, Sabino J, De Preter V, Arijs I, Eeckhaut V, et al. A decrease of the butyrate-producing species Roseburia hominis and Faecalibacterium prausnitzii defines dysbiosis in patients with ulcerative colitis. Gut. 2014;63(8):1275–83; doi: 10.1136/gutjnl-2013-304833.

18. Sokol H, Seksik P, Furet JP, Firmesse O, Nion-Larmurier I, Beaugerie L, et al. Low counts of Faecalibacterium prausnitzii in colitis microbiota. Inflamm Bowel Dis. 2009;15(8):1183–9; doi: 10.1002/ibd.20903.

19. Ferrer-Picon E, Dotti I, Corraliza AM, Mayorgas A, Esteller M, Perales JC, et al. Intestinal Inflammation Modulates the Epithelial Response to Butyrate in Patients With Inflammatory Bowel Disease. Inflamm Bowel Dis. 2020;26(1):43–55; doi: 10.1093/ibd/izz119.

20. Laserna-Mendieta EJ, Clooney AG, Carretero-Gomez JF, Moran C, Sheehan D, Nolan JA, et al. Determinants of Reduced Genetic Capacity for Butyrate Synthesis by the Gut Microbiome in Crohn’s Disease and Ulcerative Colitis. J Crohns Colitis. 2018;12(2):204–16; doi: 10.1093/ecco-jcc/jjx137.

21. Sokol H, Pigneur B, Watterlot L, Lakhdari O, Bermudez-Humaran LG, Gratadoux JJ, et al. Faecalibacterium prausnitzii is an anti-inflammatory commensal bacterium identified by gut microbiota analysis of Crohn disease patients. Proc Natl Acad Sci U S A. 2008;105(43):16731–6; doi: 10.1073/pnas.0804812105.

22. Kumari R, Ahuja V, Paul J. Fluctuations in butyrate-producing bacteria in ulcerative colitis patients of North India. World J Gastroenterol. 2013;19(22):3404–14; doi: 10.3748/wjg.v19.i22.3404.

23. Wang W, Chen L, Zhou R, Wang X, Song L, Huang S, et al. Increased proportions of Bifidobacterium and the Lactobacillus group and loss of butyrate-producing bacteria in inflammatory bowel disease. J Clin Microbiol. 2014;52(2):398–406; doi: 10.1128/JCM.01500-13.

24. Takahashi K, Nishida A, Fujimoto T, Fujii M, Shioya M, Imaeda H, et al. Reduced Abundance of Butyrate-Producing Bacteria Species in the Fecal Microbial Community in Crohn’s Disease. Digestion. 2016;93(1):59–65; doi: 10.1159/000441768.

25. Ding L, Shevach EM. IL-10 inhibits mitogen-induced T cell proliferation by selectively inhibiting macrophage costimulatory function. J Immunol. 1992;148(10):3133–9.

26. Fiorentino DF, Zlotnik A, Vieira P, Mosmann TR, Howard M, Moore KW, et al. IL-10 acts on the antigen-presenting cell to inhibit cytokine production by Th1 cells. J Immunol. 1991;146(10):3444–51.

27. Kuhn R, Lohler J, Rennick D, Rajewsky K, Muller W. Interleukin-10-deficient mice develop chronic enterocolitis. Cell. 1993;75(2):263–74; doi: 10.1016/0092-8674(93)80068-p.

28. Ostanin DV, Bao J, Koboziev I, Gray L, Robinson-Jackson SA, Kosloski-Davidson M, et al. T cell transfer model of chronic colitis: concepts, considerations, and tricks of the trade. Am J Physiol Gastrointest Liver Physiol. 2009;296(2):G135–46; doi: 10.1152/ajpgi.90462.2008.

29. Glocker EO, Frede N, Perro M, Sebire N, Elawad M, Shah N, et al. Infant colitis--it’s in the genes. Lancet. 2010;376(9748):1272; doi: 10.1016/S0140-6736(10)61008-2.

30. Glocker EO, Kotlarz D, Boztug K, Gertz EM, Schaffer AA, Noyan F, et al. Inflammatory bowel disease and mutations affecting the interleukin-10 receptor. N Engl J Med. 2009;361(21):2033–45; doi: 10.1056/NEJMoa0907206.

31. Shouval DS, Ouahed J, Biswas A, Goettel JA, Horwitz BH, Klein C, et al. Interleukin 10 receptor signaling: master regulator of intestinal mucosal homeostasis in mice and humans. Adv Immunol. 2014;122:177–210; doi: 10.1016/B978-0-12-800267-4.00005-5.

32. Kotlarz D, Beier R, Murugan D, Diestelhorst J, Jensen O, Boztug K, et al. Loss of interleukin-10 signaling and infantile inflammatory bowel disease: implications for diagnosis and therapy. Gastroenterology. 2012;143(2):347–55; doi: 10.1053/j.gastro.2012.04.045.

33. Son HJ, Kim N, Song CH, Nam RH, Choi SI, Kim JS, et al. Sex-related Alterations of Gut Microbiota in the C57BL/6 Mouse Model of Inflammatory Bowel Disease. J Cancer Prev. 2019;24(3):173–82; doi: 10.15430/JCP.2019.24.3.173.

34. Roediger WE, Truelove SC. Method of preparing isolated colonic epithelial cells (colonocytes) for metabolic studies. Gut. 1979;20(6):484–8; doi: 10.1136/gut.20.6.484.

35. Donohoe DR, Holley D, Collins LB, Montgomery SA, Whitmore AC, Hillhouse A, et al. A gnotobiotic mouse model demonstrates that dietary fiber protects against colorectal tumorigenesis in a microbiota- and butyrate-dependent manner. Cancer Discov. 2014;4(12):1387–97; doi: 10.1158/2159-8290.CD-14-0501.

36. Han A, Bennett N, MacDonald A, Johnstone M, Whelan J, Donohoe DR. Cellular metabolism and dose reveal carnitine-dependent and-independent mechanisms of butyrate oxidation in colorectal cancer cells. Journal of cellular physiology. 2016;231(8):1804–13.

37. Ribeiro WR, Vinolo MAR, Calixto LA, Ferreira CM. Use of Gas Chromatography to Quantify Short Chain Fatty Acids in the Serum, Colonic Luminal Content and Feces of mice. Bio Protoc. 2018;8(22):e3089; doi: 10.21769/BioProtoc.3089.

38. Keubler LM, Buettner M, Hager C, Bleich A. A Multihit Model: Colitis Lessons from the Interleukin-10-deficient Mouse. Inflamm Bowel Dis. 2015;21(8):1967–75; doi: 10.1097/MIB.0000000000000468.

39. De Preter V, Arijs I, Windey K, Vanhove W, Vermeire S, Schuit F, et al. Impaired butyrate oxidation in ulcerative colitis is due to decreased butyrate uptake and a defect in the oxidation pathway. Inflamm Bowel Dis. 2012;18(6):1127–36; doi: 10.1002/ibd.21894.

40. Den Hond E, Hiele M, Evenepoel P, Peeters M, Ghoos Y, Rutgeerts P. In vivo butyrate metabolism and colonic permeability in extensive ulcerative colitis. Gastroenterology. 1998;115(3):584–90; doi: 10.1016/s0016-5085(98)70137-4.

41. Roediger WE. The colonic epithelium in ulcerative colitis: an energy-deficiency disease? Lancet. 1980;2(8197):712–5; doi: 10.1016/s0140-6736(80)91934-0.

42. Amos GCA, Sergaki C, Logan A, Iriarte R, Bannaga A, Chandrapalan S, et al. Exploring how microbiome signatures change across inflammatory bowel disease conditions and disease locations. Sci Rep. 2021;11(1):18699; doi: 10.1038/s41598-021-96942-z.

43. Shinohara R, Sasaki K, Inoue J, Hoshi N, Fukuda I, Sasaki D, et al. Butyryl-CoA:acetate CoA-transferase gene associated with the genus Roseburia is decreased in the gut microbiota of Japanese patients with ulcerative colitis. Biosci Microbiota Food Health. 2019;38(4):159–63; doi: 10.12938/bmfh.18-029.

44. Vital M, Karch A, Pieper DH. Colonic Butyrate-Producing Communities in Humans: an Overview Using Omics Data. mSystems. 2017;2(6); doi: 10.1128/mSystems.00130-17.

45. Beresford-Jones BS, Forster SC, Stares MD, Notley G, Viciani E, Browne HP, et al. Functional and taxonomic comparison of mouse and human gut microbiotas using extensive culturing and metagenomics. bioRxiv. 2021:2021.02. 11.430759.

46. De Preter V, Rutgeerts P, Schuit F, Verbeke K, Arijs I. Impaired expression of genes involved in the butyrate oxidation pathway in Crohn’s disease patients. Inflamm Bowel Dis. 2013;19(3):E43–4; doi: 10.1002/ibd.22970.

47. Talero E, Alcaide A, Avila-Roman J, Garcia-Maurino S, Vendramini-Costa D, Motilva V. Expression patterns of sirtuin 1-AMPK-autophagy pathway in chronic colitis and inflammation-associated colon neoplasia in IL-10-deficient mice. Int Immunopharmacol. 2016;35:248–56; doi: 10.1016/j.intimp.2016.03.046.

48. Frank DN, St Amand AL, Feldman RA, Boedeker EC, Harpaz N, Pace NR. Molecular-phylogenetic characterization of microbial community imbalances in human inflammatory bowel diseases. Proc Natl Acad Sci U S A. 2007;104(34):13780–5; doi: 10.1073/pnas.0706625104.

49. Carlsson AH, Yakymenko O, Olivier I, Hakansson F, Postma E, Keita AV, et al. Faecalibacterium prausnitzii supernatant improves intestinal barrier function in mice DSS colitis. Scand J Gastroenterol. 2013;48(10):1136–44; doi: 10.3109/00365521.2013.828773.

50. Rossi O, Khan MT, Schwarzer M, Hudcovic T, Srutkova D, Duncan SH, et al. Faecalibacterium prausnitzii Strain HTF-F and Its Extracellular Polymeric Matrix Attenuate Clinical Parameters in DSS-Induced Colitis. PLoS One. 2015;10(4):e0123013; doi: 10.1371/journal.pone.0123013.

51. Willing BP, Dicksved J, Halfvarson J, Andersson AF, Lucio M, Zheng Z, et al. A pyrosequencing study in twins shows that gastrointestinal microbial profiles vary with inflammatory bowel disease phenotypes. Gastroenterology. 2010;139(6):1844–54 e1; doi: 10.1053/j.gastro.2010.08.049.

52. Louis P, Flint HJ. Development of a semiquantitative degenerate real-time pcr-based assay for estimation of numbers of butyryl-coenzyme A (CoA) CoA transferase genes in complex bacterial samples. Appl Environ Microbiol. 2007;73(6):2009–12; doi: 10.1128/AEM.02561-06.

53. Vital M, Penton CR, Wang Q, Young VB, Antonopoulos DA, Sogin ML, et al. A gene-targeted approach to investigate the intestinal butyrate-producing bacterial community. Microbiome. 2013;1(1):8; doi: 10.1186/2049-2618-1-8.

54. Fuentes S, Rossen NG, van der Spek MJ, Hartman JH, Huuskonen L, Korpela K, et al. Microbial shifts and signatures of long-term remission in ulcerative colitis after faecal microbiota transplantation. ISME J. 2017;11(8):1877–89; doi: 10.1038/ismej.2017.44.

55. Belzer C, Chia LW, Aalvink S, Chamlagain B, Piironen V, Knol J, et al. Microbial Metabolic Networks at the Mucus Layer Lead to Diet-Independent Butyrate and Vitamin B(12) Production by Intestinal Symbionts. mBio. 2017;8(5); doi: 10.1128/mBio.00770-17.

56. Wade H, Pan K, Duan Q, Kaluzny S, Pandey E, Fatumoju L, et al. Akkermansia muciniphila and its membrane protein ameliorates intestinal inflammatory stress and promotes epithelial wound healing via CREBH and miR-143/145. J Biomed Sci. 2023;30(1):38; doi: 10.1186/s12929-023-00935-1.

57. Png CW, Linden SK, Gilshenan KS, Zoetendal EG, McSweeney CS, Sly LI, et al. Mucolytic bacteria with increased prevalence in IBD mucosa augment in vitro utilization of mucin by other bacteria. Am J Gastroenterol. 2010;105(11):2420–8; doi: 10.1038/ajg.2010.281.

58. Swidsinski A, Dorffel Y, Loening-Baucke V, Theissig F, Ruckert JC, Ismail M, et al. Acute appendicitis is characterised by local invasion with Fusobacterium nucleatum/necrophorum. Gut. 2011;60(1):34–40; doi: 10.1136/gut.2009.191320.

59. Louis P, Duncan SH, McCrae SI, Millar J, Jackson MS, Flint HJ. Restricted distribution of the butyrate kinase pathway among butyrate-producing bacteria from the human colon. J Bacteriol. 2004;186(7):2099–106; doi: 10.1128/JB.186.7.2099-2106.2004.

60. Xie W, Gu D, Li J, Cui K, Zhang Y. Effects and action mechanisms of berberine and Rhizoma coptidis on gut microbes and obesity in high-fat diet-fed C57BL/6J mice. PLoS One. 2011;6(9):e24520; doi: 10.1371/journal.pone.0024520.

61. Hermann-Bank ML, Skovgaard K, Stockmarr A, Larsen N, Molbak L. The Gut Microbiotassay: a high-throughput qPCR approach combinable with next generation sequencing to study gut microbial diversity. BMC Genomics. 2013;14:788; doi: 10.1186/1471-2164-14-788.

62. Chen YH, Tsai WH, Wu HY, Chen CY, Yeh WL, Chen YH, et al. Probiotic Lactobacillus spp. act Against Helicobacter pylori-induced Inflammation. J Clin Med. 2019;8(1); doi: 10.3390/jcm8010090.

63. Wang L, Yang Y, Cai B, Cao P, Yang M, Chen Y. Coexpression and secretion of endoglucanase and phytase genes in Lactobacillus reuteri. Int J Mol Sci. 2014;15(7):12842–60; doi: 10.3390/ijms150712842.

64. Su J, Zhu Q, Zhao Y, Han L, Yin Y, Blachier F, et al. Dietary Supplementation With Chinese Herbal Residues or Their Fermented Products Modifies the Colonic Microbiota, Bacterial Metabolites, and Expression of Genes Related to Colon Barrier Function in Weaned Piglets. Front Microbiol. 2018;9:3181; doi: 10.3389/fmicb.2018.03181.

65. Goux HJ, Chavan D, Crum M, Kourentzi K, Willson RC. Akkermansia muciniphila as a Model Case for the Development of an Improved Quantitative RPA Microbiome Assay. Front Cell Infect Microbiol. 2018;8:237; doi: 10.3389/fcimb.2018.00237.

