## Supplemental figure for "Inflammation is the Driver of Butyrate-Producing Bacteria Change in Interleukin10 Knockout Mice"

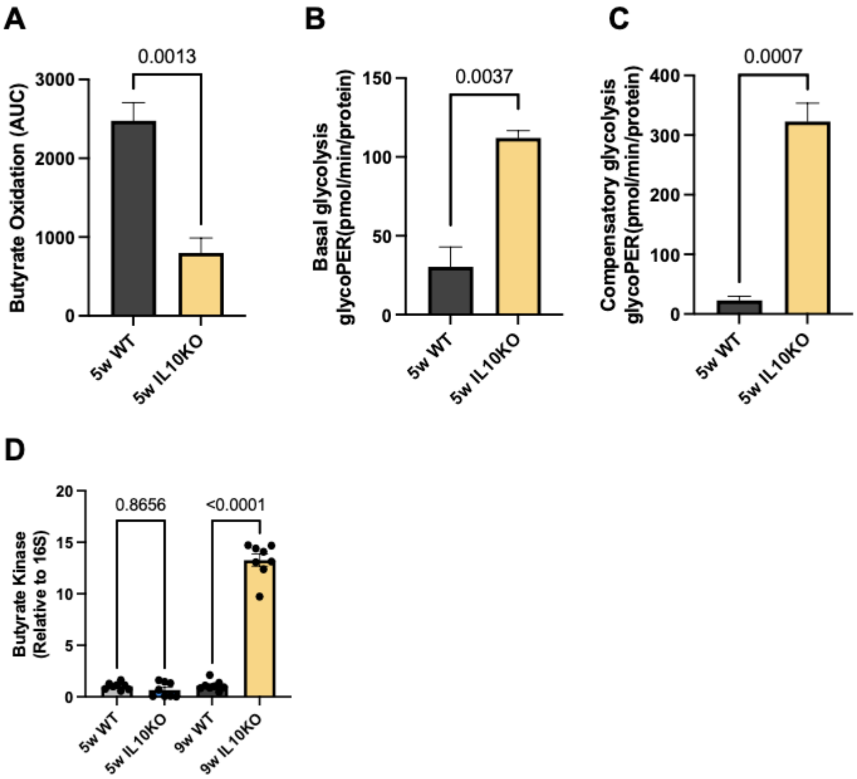


**Supplemental Figure S1. Response of 5-week-old IL10 Knockout Mice to Butyrate and Glucose, and Lower Butyrate Kinase Levels**

**(A)** Butyrate oxidation represents the area under the curve (AUC) from oxygen consumption rate (OCR) measurements taken after 2DG injection but before sodium azide (SA) injection. Data points represent the OCR per condition for butyrate oxidation measurements. Contribution of basal glycolysis **(B)**, and compensatory glycolysis **(C)** in wildtype (WT) mice and IL10 knockout (IL10 KO) mice. Data points represent the glycoPER per condition for glycolysis measurements. **(D)** Abundance of butyrate kinase (buk) was measured in the fecal colonic contents of 5-week-old and 9-week-old WT mice and IL10 KO mice. Error bars indicate the mean ± SD (n = 4,5, or 7). Significant differences are shown.

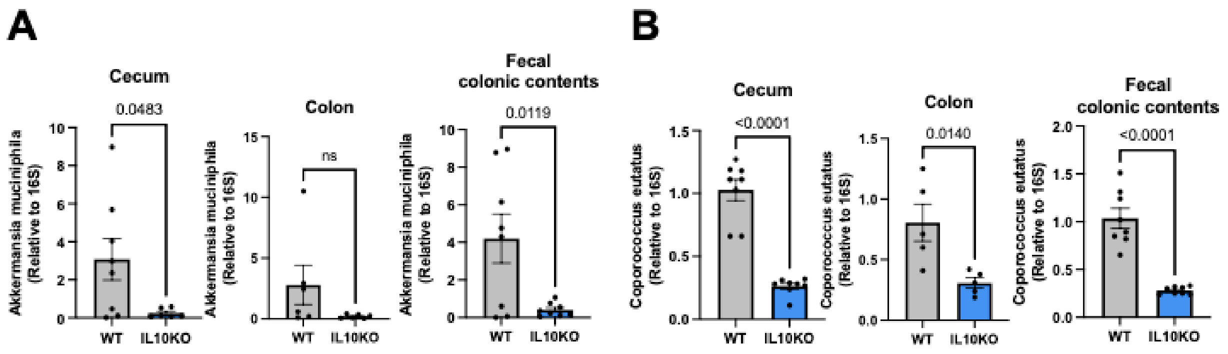


**Supplemental Figure S2. Abundance of *Akkermansia muciniphila* and *Coprococcus eutatus* at the Species level in IL10 Knockout Mice**

**(A)** Abundance of the species *Akkermansia muciniphila* was measured in the cecum, colon, and fecal colonic contents of wildtype (WT) mice and IL10 knockout (IL10 KO) mice. **(B)** Abundance of the species Coprococcus eutatus was measured in the cecum, colon, and fecal colonic contents of WT mice and IL10 KO mice. Error bars indicate the mean ± SD (n = 6 or 8). Significant differences are shown.
